# A delay-deterministic model for inferring fitness effects from time-resolved genome sequence data

**DOI:** 10.1101/229963

**Authors:** Nuno R. Nené, Alistair S. Dunham, Christopher J. R. Illingworth

## Abstract

A common challenge arising from the observation of an evolutionary system over time is to infer the magnitude of selection acting upon a specific genetic variant, or variants, within the population. The inference of selection may be confounded by the effects of genetic drift in a system, leading to the development of inference procedures to account for these effects. However, recent work has suggested that deterministic models of evolution may be effective in capturing the effects of selection even under complex models of demography, suggesting the more general application of deterministic approaches to inference. Responding to this literature, we here note a case in which a deterministic model of evolution may give highly misleading inferences, resulting from the non-deterministic properties of mutation in a finite population. We propose an alternative approach which corrects for this error, which we denote the delay-deterministic model. Applying our model to a simple evolutionary system we demonstrate its performance in quantifying the extent of selection acting within that system. We further consider the application of our model to sequence data from an evolutionary experiment. We outline scenarios in which our model may produce improved results for the inference of selection, noting that such situations can be easily identified via the use of a regular deterministic model.

## Introduction

Fitness landscapes describe the relationship between the genome of an organism and its evolutionary fitness^1^. Evolutionary fitness encompasses a broad range of important phenotypes of an organism, making the inference of details of fitness landscapes a topic of broad biological interest. In some important biological systems, adaptation occurs as a rapid and ongoing process^2,3^. Where multiple beneficial mutations arise in a population simultaneously, linkage between mutations has a substantial impact upon adaptation; a considerable body of literature has characterised the implications of such effects for adaptation^4–11^.

Where adaptation is sufficiently rapid to be observed, time-resolved sequence data may be of assistance in measuring the extent to which a variant is under selection. Under the assumption of a large population size, the evolution of a single beneficial allele over time can be described by deterministic differential equations^12^. Given sufficient observations of a population under study, the simplicity of this deterministic framework allows it to be extended to infer selection in far more complicated evolutionary scenarios^13,14^; fitting a deterministic model to data provides an estimate of the magnitude of selection acting upon one or very many alleles. In other situations, genetic drift is an important factor to account for; in a small population, changes in allele frequency occurring via drift may outweigh those caused by selection^15^. In this situation, a variety of methods have therefore been developed to consider the evolution of a single-locus, two-allele system, estimating in a joint calculation the effective size of a population, and the magnitude of selection acting upon a variant allele^16–23^. In a similar calculation, one may estimate whether or not a change in the frequency of an allele has arisen through selection or genetic drift. Genetic drift induces an uncertainty in the future frequency of an allele^24^; accounting for this, alleles which have changed by more than a given threshold may be identified, enabling the attribution of selection to genetic variants^25–27^. A similar approach has been applied to the case where a population is large, but measurements of allele frequency are uncertain; model selection procedures discriminate ‘neutral’ from ‘selected’ behaviour in an allele frequency trajectory^28^.

Where genetic drift is incorporated into a model, a variety of approaches to modelling Wright-Fisher propagation have been adopted^29^. Numerical solution of the stochastic dynamics of the population may be computationally intensive, inspiring the development of more rapid propagation methods, and the consideration of potential alternative solutions^30–32^. In a recent work, examining a range of potential models for the demographic history of a population, it was concluded that deterministic approximations to evolution under drift can produce accurate estimates of the magnitude of selection^33^. Such models of selection, mutation, and recombination have been used to generate insights into viral adaptation^34–36^. Time-resolved sequence data describing pathogenic populations is becoming increasingly available^37–40^; in so far as demographic effects can be ignored in such systems, evolutionary inference becomes possible at far-reduced computational cost, making this an important area for methodological development and application.

While acknowledging the potential for deterministic models to generate biological insight, we here present an important case in which a deterministic inference of selection from population sequence data produces a severely deficient result. In this case, a stochastic approach to inference produces a correct result, albeit with additional prior knowledge of the system and at the cost of a substantial amount of computational time. However, the use of what we term a delay-deterministic model, including a single extra model parameter, goes a substantial way to correcting the error in the deterministic calculation. We propose that under a range of circumstances we go on to describe, the delay-deterministic model combines the speed of a deterministic modelling framework with the accuracy provided by more computationally intensive models.

## 1 Materials and Methods

### 1.1 Simulated trajectories under a Wright-Fisher propagation model

Simulated data from a population was generated according to a model of sequential mutation and selection steps. We consider a population of *N* individuals occupying a linear network of *L* + 1 distinct haplotypes, each haplotype being separated from the previous one by a single mutation. We model the fitness of each haplotype as continually increasing with linear distance, such that the fitness of haplotype *i* is given by *w_i_* = 1 + *si* for some parameter *s* (Figure 1). We denote the number of individuals of haplotype *i* in the population after *t* generations by *n_i_*(*t*). Within each generation, propagation of the system was conducted using a simple model of mutation and selection. Mutation was modelled as occurring between adjacent haplotypes; for a given mutation rate *μ*, the number of mutants *m_ij_* from haplotype *i* into an adjacent haplotype *j* was given by the Poisson distribution

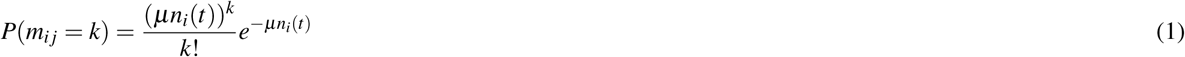

Subsequently, the next generation was drawn via a multinomial process

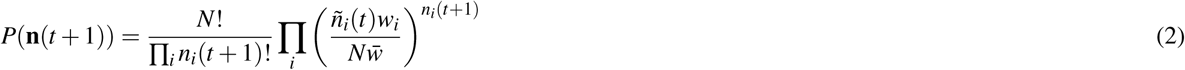

where *w_i_* is the fitness of the haplotype *i*, *ñ_i_*(*t*) is the number of individuals of haplotype *i* at time *t* after the effect of mutation has been accounted for, and *w̅* is the mean fitness of the population. Sequencing of the population was simulated via a multinomial emission model with sequencing read depth *N_d_*. We consider the behaviour of our system with a variety of parameters. Simulations were conducted with *μ* = 10^−5^, population size *N* ∈ {10^5^,10^9^}, *s* ∈ {0.1,0.2,…, 0.9}. A sequencing depth of *N_d_* = 10^3^ was simulated, with sampling from the population every generation; such a depth of sampling is likely unrealistic for a biological system but allows for clearer comparison of different inference methods. Our model therefore simulates the effect of strong selection, with *Ns* ≫ 1, but is not restricted to the strong mutation paradigm of *μ N* ≫ 1^15,41^. In each case the initial state of the system was defined by *n*_0_(0) = *N* and *n_i_*(0) = 0 for all *i* > 0.

**Figure 1.**
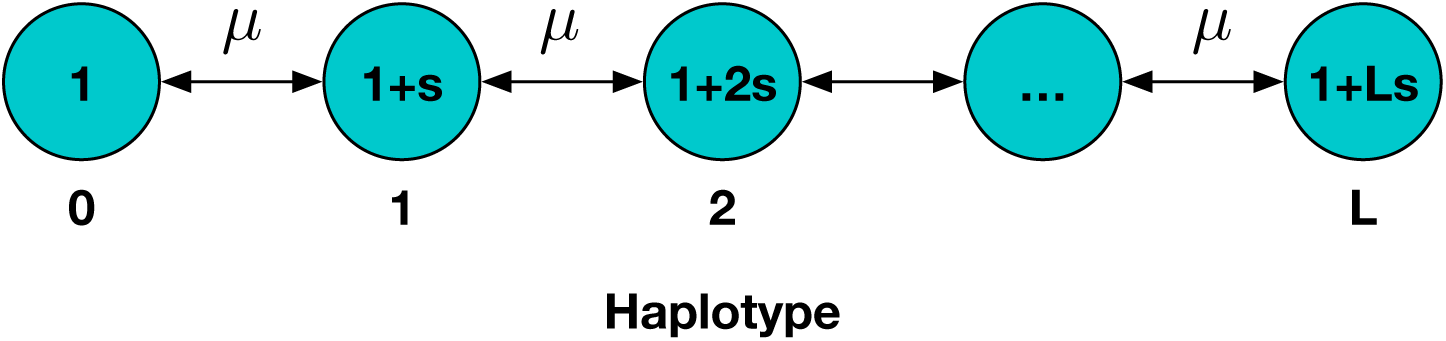
Linear haplotype model used for simulation. The *i^th^* haplotype has fitness *w_i_* = 1 + *is*; mutation occurs between adjacent haplotypes with constant rate *μ*.

### 1.2 Inference methods

Inferences of selection were conducted using an evolutionary model to generate inferred haplotype frequencies *q_i_*(*t*) across time. Given a set of such frequencies, a multinomial log likelhood was calculated for the system

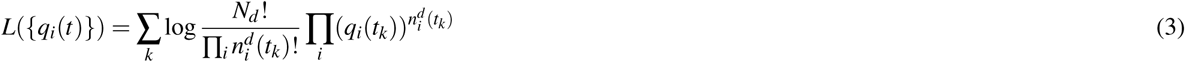

where 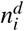 (*t_k_*) is the number of observations of the haplotype *i* at time *t*_k_, and the sum is calculated over data from all observed time points. The parameters *q_i_*(*t*_0_) and *s* were optimised in each case. Three models were used to generate inferred frequencies.

#### 1.2.1 Deterministic inference model

In the first model, haplotype frequencies were modelled under the assumption of an infinite population size. As such, in each generation a fraction of each haplotype was subject to mutation

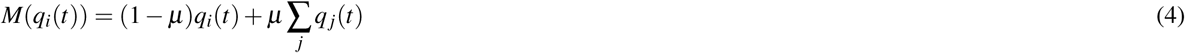

where as in the simulations the sum is conducted over all haplotypes *j* that differ from *i* by a single allele. Selection was included in a similarly deterministic manner:

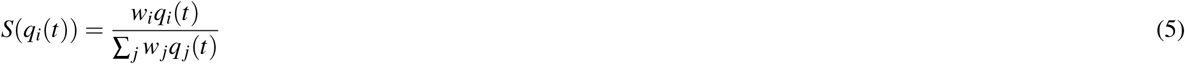

where the sum was calculated over all haplotypes. The next generation is given by

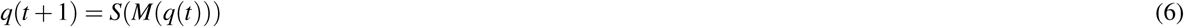

#### 1.2.2 Stochastic inference model

In the second model, allele frequencies were propagated in exactly the same way as in the model used for simulation. Stochastic simulations of viral populations have been used to explore the potential range of outcomes occurring in viral systems^42^. In order to sample the space of potential outcomes of the model, 1000 replicates of the model were run for each proposed set of parameters; the mean value of the likelihoods for the replicates, each calculated according to Eq.3, was maximised in order to identify parameters. In order to deal with the stochasticity of the likelihood function, an optimisation method was implemented that prevented the resampling of previously-tested model parameters.

#### 1.2.3 Delay-deterministic model

Finally a delay-deterministic model was implemented, identical to the deterministic model described above, but with the addition of a delay representing the time for establishment of individuals with a novel haplotype. Specifically, the mutation function of Eq.4 was modified, with mutation out of a haplotype occurring only if the frequency of that haplotype was greater than a specific threshold.

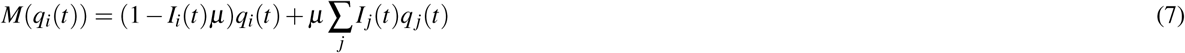

where the index function

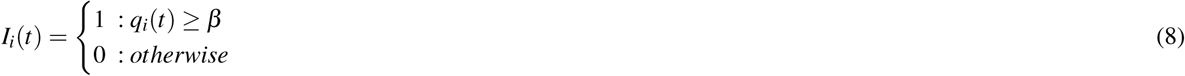

The parameter *β* was optimised to identify the maximum likelihood model.

We note that, of the three inference models, the stochastic model requires an estimation of the total population size, *N*; for the sake of computational time we used the correct value of this parameter in our inferences. Neither the deterministic or delay-deterministic models require an estimate of population size.

#### 1.2.4 Application to experimental data

To explore the use of our approach with experimental data, the deterministic and delay-deterministic models were applied to influenza sequence data collected from an evolutionary experiment conducted in ferrets^43^. A previous analysis of these data using a deterministic approach has been described in an earlier publication^35^; for simplicity we here considered within-host data from a single animal in the study, denoted F3501 in the original work, for which the initial population diversity was relatively low. Genome sequence data were processed as described in the previous publication^35^, identifying loci in the HA segment at which significant change in allele frequency was observed, then processing short read data spanning these loci into a set of multi-locus variant calls and inferring haplotype frequencies which best fit the observed data using a maximum likelihood model. Mutation was initially modelled as occurring deterministically between haplotypes, identifying an optimal model of haplotype fitness using model selection. Fitness parameters within this model were then re-inferred using the delay-deterministic framework, comparing inferred fitnesses from under each approach. In common with the default model in the original study, the rate of mutation was modelled as *μ* = 10^−5^.

## 2 Results

The stochastic and delay-deterministic models produced the most accurate inferences of selection. Selection coefficients inferred from the three different models are shown in Figure 2. For each of the values of *μN*, the deterministic model underestimates the magnitude of selection, *s*, for values of *L* greater than one (that is, where there were three or more haplotypes in total), with an increasing degree of underestimation as *L* increases. Where *L* = 5, the gradient of a linear model fitted to the mean inferred frequencies was equal to 0.095; roughly one tenth of what would be given by a correct inference model. The results obtained also depend upon the value of *μN*; where this statistic is larger, the extent to which the deterministic model underestimated the true magnitude of selection was reduced; here for the case *L* = 5 the gradient of the fitted linear model was 0.39. Results from the stochastic inference model show/ a good reproduction of the correct fitness values; linear gradients varied between 0. 98 and 1.02 for the case *μN* = 1, indicating an accurate reproduction of selection coefficients as would be expected given the identical models of propagation. The delay-deterministic method was close in performance to the stochastic model, with gradients between 0.88 and 0.97 indicating a small underestimate of the strength of selection. This underestimate was not reproduced in the case *μN* = 10000, for which gradients fitted to the delay-deterministic outputs were either side of 1.

**Figure 2.**
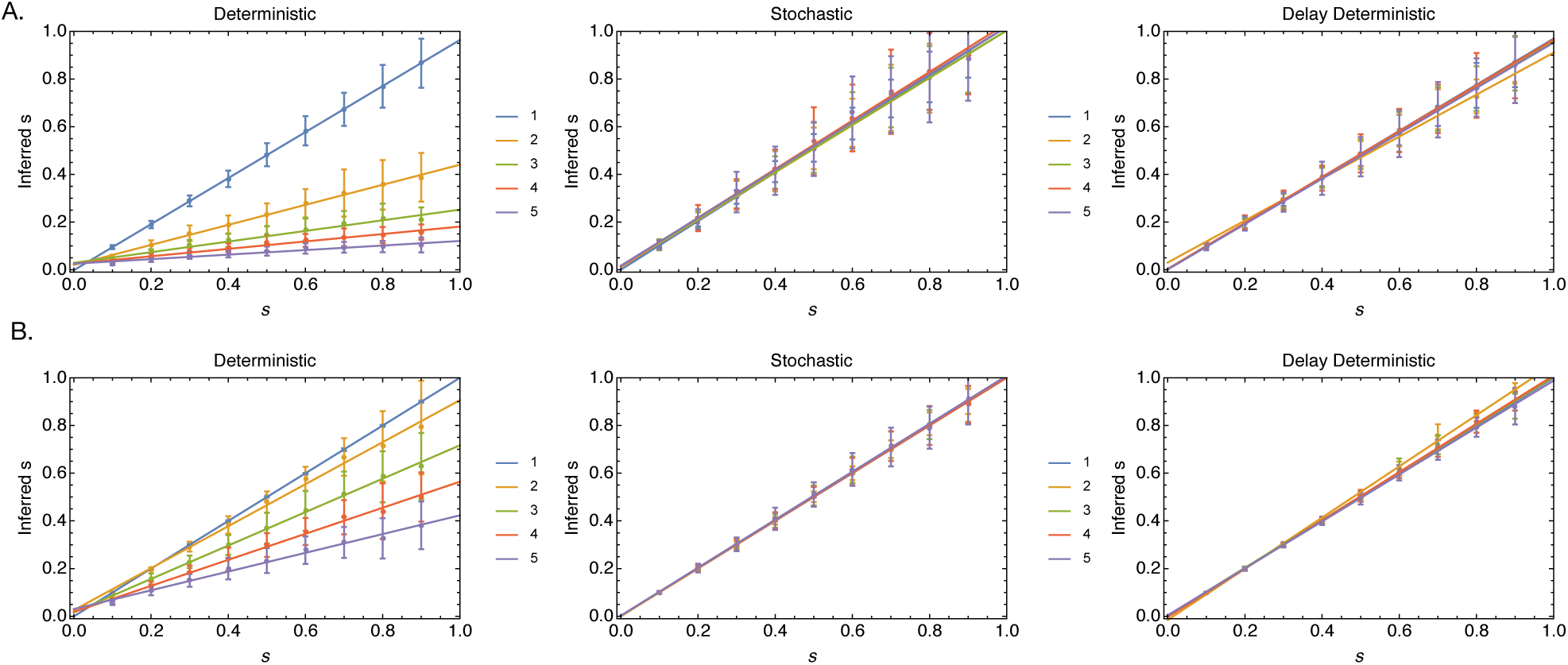
The deterministic model substantially underestimates the correct magnitude of selection for models with multiple haplotypes. Parameters are shown for values of L between 1 and 5 with (A) *μN* = 1. (B) *μN* = 10000. Points show mean inferred selection coefficients; error bars were determined from a set of 100 replicate calculations. Lines show the outputs of a linear regression model fitted to the mean values.

The results we obtained can be intuitively understood via a plot of the evolutionary dynamics of the linear system (Figure 3). Given a deterministic model with the correct selection coefficient, the population propagates through the haplotypes substantially faster than does the stochastic model. In the Wright-Fisher model, given that *Nμ* = 1, a mean of one individual mutates from haplotype 0 to haplotype 1 in the first generation. Following the second generation the probability of an individual being found in haplotype 2 is therefore approximately *μ*. In so far as double mutations are ignored within our model framework, at least one individual is required to occupy a haplotype before the next haplotype can be founded via mutation; this leads to a delay of multiple generations before a single individual reaches the final haplotype, at which point selection of magnitude of 1 + *Ls* ensures the eventual fixation of this haplotype. By contrast, in the deterministic model, mutation propagates the population rapidly through the system; after L generations the final haplotype is deterministically occupied by a frequency of the population of order *μ^L^*. The increased fitness of this final haplotype therefore takes effect on the system more rapidly, leading to the observed faster propagation. When the deterministic model is optimised, a lower fitness parameter s is inferred to compensate for this effect, which increases dependent upon the number of haplotypes in the system. By contrast, the delay-deterministic model corrects for the error of the deterministic model. By imposing a delay on the rate at which new haplotypes are founded by mutation, the rate of propagation through the haplotypes is controlled, giving an improved fit to the data, and therefore a more accurate inference of selection.

**Figure 3.**
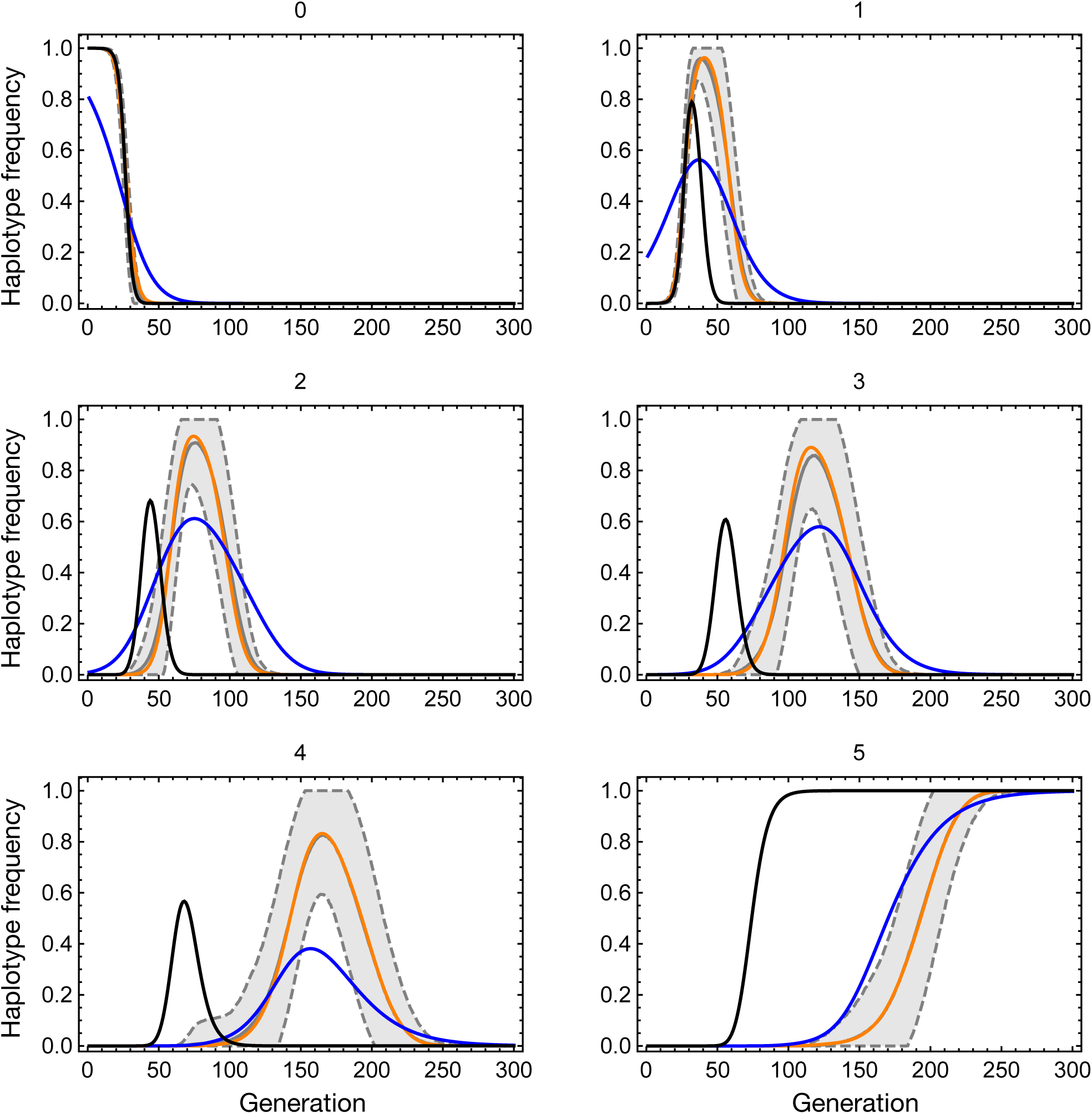
Time-dependent dispersion of trajectories for the case *μN* = 1, *L* = 5 and *s* = 0.5. Frequencies of each haplotype are shown reading left to right from the top. In each case the solid gray line (sometimes obscured) shows the mean haplotype frequency of the simulated data across time, calculated across 100 simulations. The region within one standard deviation of this frequency is indicated by gray dashed lines and is shaded. The black line shows the propagation of the deterministic model in the case where *s* = 0.5 and the population starts at the same haplotype distribution as the simulation, while the blue line shows the results of a maximum likelihood fit between the deterministic model and the mean data. The mean maximum likelihood fit of the delay-deterministic model to the data is shown as an orange line.

At higher values of *μN*, differences between the stochastic and deterministic systems are reduced (Supporting Figure S1).

As *N* tends to infinity, the number of individuals mutating between haplotypes per generation approaches the deterministic limit and the time at which a haplotype becomes established decreases, with the consequence that less of a reduction in *s* is required to fit the model to the data. The deterministic model therefore provides a good description of the behaviour of the discrete system as *μ^L^N* in the discrete model approaches a value much larger than 1. For a within-host model of influenza, where *μ* may be of the order 10^−44^, and N potentially of the order 10^14^^42^, this implies that the a value of *L* of 4 or greater could lead to failure of the deterministic model; we next consider the application of our models to data from an experimental evolution study.

Application of the deterministic and delay-deterministic methods to data from an evolutionary experiment^43^ showed only small differences between inferred parameters. Details of each inference are given in Supporting Table S1. Within this system evolution proceeds exceptionally fast, with mutation into new and highly advantageous haplotypes being inferred to drive the adaptation of the system over the course of an infection (Figures 4A, 4B). Application of the delay-deterministic value gave a marginally improved fit to the data, with a maximum likelihood value 0.63 units better than the deterministic model without accounting for the additional parameter. The value of *β* was inferred to be 1.8 × 10^−10^, smaller than any inferred initial haplotype frequency that was greater than zero. However, the inferred haplotype fitness values were very similar between the models, with deviations in fitness of not much greater than 1% (Figure 4C). This final result can be understood in terms of the arrangement of haplotypes within the system; although some haplotypes were inferred to have initially zero frequency, being created by mutation from other haplotypes, there was insufficient time for more distant haplotypes, at least two mutations away from initially occupied haplotypes, to increase to an appreciable frequency. This result is informative for calculations performed on biological datasets; even where selection for novel variants is extreme in nature, a delay-deterministic model is unlikely to be required to generate correct inferences of selection on timescales of 4-5 days. Previous inferences of selection for within-host influenza adaptation using deterministic methods are therefore unlikely to be negatively affected by the use of a deterministic model of mutation^36^. Rather, the value of the method will arise over longer timescales, where the population grows under selection into haplotypes that are separated by multiple variants from sequences in the initial population.

**Figure 4.**
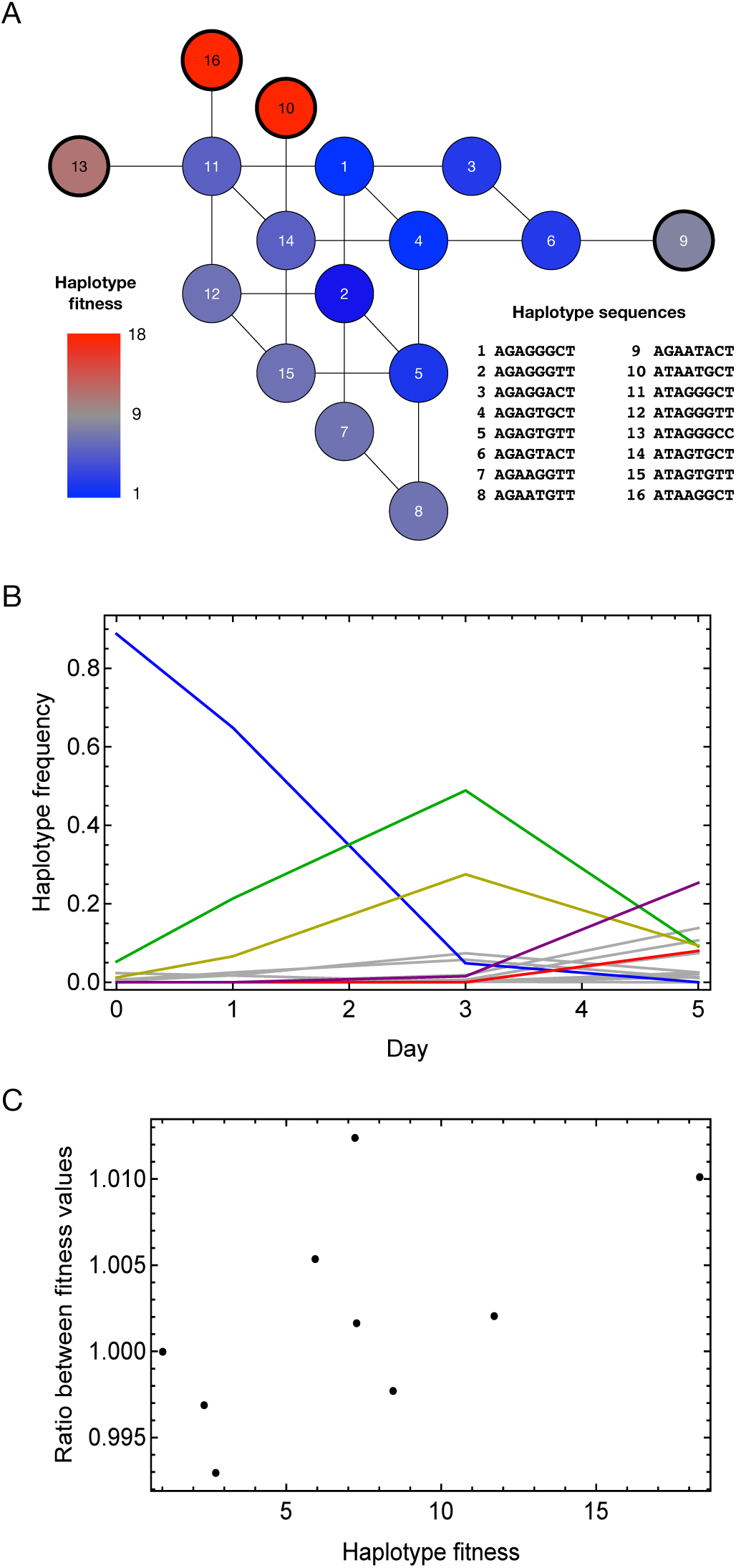
(A) Fitness landscape inferred from experimental data using the deterministic method. Inferred fitnesses are given for haplotypes for which the inferred frequency reached at least 1% within the course of within-host evolution. Given haplotypes describe the sequence of the HA segment of the virus at genome positions 339, 496, 728, 738, 788, 1018, 1020 and 1144. The haplotypes 9, 10, 13, and 15, marked in darker outline, were inferred to have zero initial frequencies. (B) Inferred changes in haplotype frequency over time. Haplotype frequencies are shown in gray, with the exceptions of haplotypes 1 (blue), 2 (green), 6 (yellow), 12 (purple) and 16 (red). (C) Differences in inferred haplotype fitness values between the deterministic and delay-deterministic methods shown proportional to the value inferred under the deterministic model.

## 3 Discussion

Deterministic models of adaptation have been proposed as a rapid and effective method for inferring selection from time-resolved sequence data^33^. We have here highlighted a limitation of such frameworks whereby a deterministic model may severely underestimate the magnitude of selection in a system. This underestimation results from delays in the propagation of a finite population towards mutationally distant haplotypes; at least one individual is required to occupy a haplotype before mutation out of that haplotype may occur. As shown here, this delay operates even at high values of *Nμ*; we suggest that a population size satisfying *Nμ^L^* ≫ 1 would be required to remove this effect. As a solution to this problem we propose an alternative inference procedure, which we term a delay-deterministic model. Under this model the progress of a nominally infinite population through the system is delayed via the use of an additional model parameter, matching the behaviour of the stochastic system; the additional parameter recreates the behaviour of a finite population, giving a correct inference of selection.

In demonstrating the application of our model, we have chosen the simplest possible situation in which the effects we are studying apply; that of a linear set of haplotypes separated by single mutations. Such a system has previously been considered in an application to cancer, calculating the time at which a novel haplotype might arise^45^; while the system we consider is similar, our research question differs from this earlier study. We note that our approach is not limited to the linear set in so far as it can be applied to the inference of fitness landscapes of greater complexity. We further note that our model is not the only approach that would give a correct inference of selection under the circumstances of a population entering mutationally distant haplotypes. For example, inferring an ‘establishment time’ for each haplotype, at which a haplotype enters a population at a frequency above the selection-drift threshold^46^, would likely generate correct results, albeit at the cost of learning a single parameter per haplotype in the system. The use of a model of time-dependent selection, in which selection only begins to affect a haplotype at a specific point in time^47^ would also give an approximately correct inference of selection, delaying the impact of selection to a point at which the inferred trajectory with fit the model. However, this again would incur a computational cost, and would require the imposition on the system of the potentially incorrect assumption of a change in the magnitude of selection with time. Considering approximations to the stochastic system an approach based upon a reproduction of the stochastic distribution of allele frequencies^48^ would likely give a correct result albeit this has not to our knowledge been achieved. The delay-deterministic model we present here gives a computationally rapid approach to correctly infer the magnitude of selection.

While accurately reproducing the magnitude of selection, our model has an advantage over the stochastic approach in utilising a framework of deterministic propagation. Whereas, in the manner we implemented it, the stochastic model required 1000 replicate propagations of the model for each set of parameters tested, the delay-deterministic approach requires only the optimisation of a single additional parameter. The relative cost of this is likely to vary considerably depending upon the complexity of the system in question; in the application to influenza data considered here, where the model contained tens of parameters to be optimised, a single additional parameter is not likely to add substantially to the computational cost, provided the optimisation procedure is implemented in an efficient manner. We note that faster implementations of the stochastic framework are likely to be achievable; the recent demonstration of such methods for the two-allele case suggest the extension to more generalised population models to be a valuable avenue for exploration^30,32^.

Applying our model to data from an evolutionary experiment, we identified very similar results between the deterministic and delay-deterministic methods; despite very high magnitudes of selection being inferred to act upon haplotypes in this case, little difference in the model inferences was found. We therefore propose that our method will be of relevance in cases where selection is strong and acts over longer time periods than those of the experiment considered, for which adaptation was observed over only a 5-day window. The identification of cases for which the deterministic model will produce correct or incorrect results is likely to be possible via application of the model itself; wherever a substantial proportion of a population is inferred to evolve into a haplotype more than two mutations distant from a haplotype occupied by the initial population, a delay-deterministic or similar approach should be considered.

